# Bone morphogenetic protein 4 reduces global H3K4me3 to inhibit proliferation and promote differentiation of human neural stem cells

**DOI:** 10.1101/2020.01.22.915934

**Authors:** Sonali Nayak, Benjamin Best, Emily Hayes, Ashorne Mahenthiran, Nitin R Wadhwani, Barbara Mania-Farnell, Rintaro Hashizume, John A Kessler, Charles David James, Tadanori Tomita, Guifa Xi

## Abstract

Posttranslational modifications (PTMs) on histone tails spatiotemporally dictate mammalian neural stem cell (NSC) fate. Bone morphogenetic protein 4 (BMP4), a member of the transforming growth factor β (TGF-β) superfamily, suppresses NSC proliferation and fosters differentiation into astroglial cells. Whether PTMs mediate these effects of BMP4 is unknown. Here we demonstrate that BMP4 signaling causes a net reduction in cellular histone H3 lysine 4 trimethylation (H3K4me3), an active histone mark at promoters of genes associated with human NSC proliferation. We also show that H3K4me3 reduction by BMP4 is mediated by decreased expression of SETD1A and WDR82, two methyltransferase components of SETD1A-COMPASS. Down-regulation of these components decreases expression of key genes expressed in hNSCs, while ectopic expression via transfection dedifferentiates human astrocytes (HAs). These observations suggest that BMP4 influences NSC fate by regulating PTMs and altering chromatin structure.

**SIGNIFICANCE STATEMENT:** BMP4 is critical in determining hNSC fate. Whether histone posttranslational modifications (PTM) mediate the effects of BMP4 is unknown. Here we report that H3K4me3, brought about by its methyltransferases SETD1A and WDR82, at promoters of stem cell genes OCT4 and NESTIN is involved in human neural stem cell (hNSC) maintenance. BMP4 promotes hNSC astroglial differentiation in part through reduction of SETD1A and WDR82 and thus decreased frequency of H3K4me3 at the promoters of these genes. These results provide evidence that BMP4 promotes hNSC differentiation through a potential epigenetic mechanism and extend our understanding of the role of histone PTM in central nervous system development.

## INTRODUCTION

Neural stem cells (NSCs) either self-renew or differentiate into neurons, astrocytes and oligodendrocytes [1, 2]. Higher order chromatin structure precisely regulates gene expression to dictate NSC fate [3, 4]. Histones are key components of chromatin and subject to a wide variety of dynamic post-translational modifications (PTMs) [5]. Histone and associated modifications are important determinants of human NSC self-renewal, proliferation and differentiation [4, 6-9].

Trimethylation of histone H3 lysine 4 (H3K4me3) by a complex of proteins associated with SET1A/B (SETD1A/B-COMPASS) [10-12], is pivotal in determining human NSC fate [13-17]. H3K4me3 is enriched near transcription start sites (TSSs) at the 5’ regions of virtually all actively-expressed genes; its presence has a strong positive correlation with gene transcription rates, polymerase II promoter occupancy [18, 19], and transcriptional consistency [20]. WDR82 is a specific subunit of SETD1A/B. Other subunits of SETD1A/B-COMPASS include WDR5 [21], DPY30 [22], and ASH2L [23].

Bone morphogenetic protein 4 (BMP4), a member of the TGF-β superfamily, is essential for central nervous system (CNS) development due to its influence on NSC self-renewal, proliferation and differentiation [24]. BMP4 signaling affects the expression of key genes that affect NSC fate. For example, BMP4 increases CXXC finger protein 5 (CXXC5) to repress Wnt signaling and promote NSC differentiation [25]. It increases expression of inhibitor of differentiation 4 (ID4) and ID2, which inhibits oligodendroglial lineage commitment and promotes astrocyte commitment [26]. BMP4 inhibits mitogen-activated protein kinase (MAPK) pathways that support stem cell self-renewal [27], and it stimulates msx2 and p21 to mediate pro-apoptotic effects on NPCs in the ventricular zone [28].

However, little is known regarding effects of BMP4 on chromatin structure and how these changes influence gene expression. BMP4 differentiates CD133-positive glioma stem cells to astroglial-like cells [29] and inhibits their proliferation through decreasing CYCLIN D1 expression [30]. SMAD-mediated BMP4 signaling regulates embryonic stem cell proliferation by altering H3K4me3 enrichment at promoters of proliferation genes [31]. In NSCs, the CYCLIN D1 promoter is enriched for H3K4me3 [20]. We found that Human StemPro® NSCs are CD133 positive which led to the hypothesis that BMP4 inhibits human StemPro® NSC proliferation through reduction of H3K4me3 at the CYCLIN D1 gene. In this study we show that BMP4 signaling regulates H3K4me3 and chromatin structure, thereby influencing expression of several genes with important consequences for NSC differentiation and proliferation.

## MATERIAL AND METHODS

### Reagents and antibodies

Knockout™DMEM/F-12(Cat#12660-012), StemPro® neural supplement (Cat#A10508-01), epidermal growth factor (EGF), fibroblast growth factor-2 (FGF-2), GlutaMAX™-I CTS™ (100X) (Cat#A12860), Geltrex® hESC-qualified Reduced Growth Factor Basement Membrane Matrix (Cat# A1413301), StemPro® Accutase® Cell Dissociation Reagent (Cat#A11105), and Geltrex® LDEV-Free hESC-qualified Reduced Growth Factor Basement Membrane Matrix (Cat#A14133) were purchased from Life Technologies (Frederick, MD, USA). Accutase was purchased from Millipore (Billerica, MA, USA). Heparin (Cat#H3149) and ascorbic acid (Cat#A8960) were from Sigma Aldrich (St. Louis, MO, USA).

Rabbit polyclonal antibody against WDR82 was a generous gift from Dr. David G. Skalnik (Indiana University School of Medicine, IN, USA). Antibodies were purchased from the following companies: BMP4 (sc-6896), OCT4 (sc-8629), and GAPDH (sc-25778) from Santa Cruz Biotechnology (Dallas, TX, USA); human SETD1A (Cat#A300-289A) and SETD1B (Cat#302-280A) from Bethyl Laboratories (Montgomery, TX, USA); H3K4me3 (Cat#9727), Histone H3 (Cat#9715), ASH2L (Cat#5019); WDR5 (Cat#13105), RBBP5 (Cat#13171), and CYCLIN D1 (Cat#2922) from Cell Signaling Technology; CD133 (ab5558), and β-actin (ab-8227) from Abcam (Cambridge, MA, USA).

### Immunohistochemistry

Formalin-fixed, paraffin-embedded (FFPE) fetal brain tissue slides containing lateral ventricle lining from autopsy brain samples, from two pediatric patients who died of non-brain related diseases, were obtained from the Department of Pathology, Ann & Robert H. Lurie Children’s Hospital of Chicago. FFPE slides were de-paraffinized in xylene, and hydrated through a graded series of alcohols. Endogenous peroxidases were blocked with 3% hydrogen peroxide. Antigen retrieval was performed by boiling for 20 min in a 0.01 M sodium citrate (pH 6.0) solution and endogenous biotin was blocked using the Avidin/Biotin Blocking Kit (Vector Labs, SP-2001). Slides were incubated overnight with primary antibodies against anti-rabbit H3K4me3 (Cell Signaling Technology, Cat#9727, 1:400). Following incubation with biotin-labeled secondary antibodies (Vector Laboratories BA1000), antigens were visualized with streptavidin-biotin (Vectastain Elite ABC kit; Vector Laboratories) followed by Vector^®^ NovaRED™ (Vector Laboratories) and counterstained with hematoxylin (Richard-Allen Scientific). Images were captured on a Leica DMR-HC upright microscope and analyzed using OpenLab 5.0 software.

### Cell culture: primary neurospheres, spontaneously differentiated cells and human astrocytes

Human StemPro® NSC line derived from fetal brain was purchased from Life Technologies, Inc. (Grand Island, NY, USA). For neurospheres, the cells were cultured in StemPro® NSC serum free medium (SFM), prepared with Knockout™ DMEM/F-12 medium including 2% StemPro® Neural Supplement, 20ng/ml bFGF, 20ng/ml EGF, 2mM GlutaMAX™-I supplement, 6 units/ml heparin and 200μM ascorbic acid. For routine passaging, cells were dissociated with StemPro® Accutase® Cell Dissociation Reagent. For spontaneous differentiation, cells were harvested as single cell suspensions with StemPro® Accutase® Cell Dissociation Reagent as per manufacturer’s instruction. The cells were re-suspended in differentiation medium prior to plating on Geltrex® matrix-coated culture vessels, and grown with SFM without bFGF and EGF. The medium was changed after 2 days followed by replacement of half of the media every 2–3 days. Following 7 days of differentiation, all three neural lineages (astrocytes, neurons, and oligodendrocytes) were present. The normal human astrocyte cell line (HA) was kindly provided by Dr. Rintaro Hashizume and grown as described [32, 33].

### Immunofluorescence

Seven days after primary neurospheres culture, cells were harvested as single cells and split with SFM or differentiation medium. For sphere staining, spheres were plated onto glass coverslips coated with Geltrex® matrix placed in individual wells of a 12-well culture plate in SFM and monitored under a microscope every 15mins. Once spheres attached and began to differentiate they were fixed with 4% paraformaldehyde in PBS (Pierce Chemical Co., Rockford, IL). For staining of differentiated cells, cells were plated onto glass coverslips coated with Geltrex® matrix, and cultured in individual wells of a 12-well culture plate with SF differentiation medium. Seven days after plating, coverslips were fixed with 4% paraformaldehyde in PBS. Fixed cells were blocked with 10% donkey serum and 0.3% Triton X-100 in PBS and incubated with CD133 (1:100), OCT3/4 (1:100), SOX2 (1:100), and NESTIN (1:100) for neurospheres; and with MUSHASHI (1:100), GFAP (1:200), β-TUBULIN (1:100) for neurons, and O4 (1:100) for differentiated astrocytes, neurons and oligodendroglial cells.. Alexa Fluor 488 or cy3 labeled secondary antibodies (dilution 1:200) (Jackson Lab, ME, USA) were used for detection. Nuclei were counterstained with DAPI. Images were captured with a Leica DM-IRB inverted microscope and analyzed using OpenLab 5.0 software.

### Constructs and transfections

pcDNA3-WDR82-HA was a generous gift from Dr. David G. Skalnik (Indiana University School of Medicine, IN, USA). pcDNA3-HA was a gift from Kunliang Guan (Addgene plasmid #13512). Human His-tagged SETD1A expression pET28-SETD1A-MHL plasmid and its control pET28-MHL were gifts from Cheryl Arrowsmith (Addgene plasmid # 32868 and #26096, respectively). Human short interference RNA (siRNA) against human SETD1A (Gene ID 9739, Cat# SR306505), short hairpin RNA (shRNA) against WDR82 (Gene ID 80335, Cat#TG301034), and scrambled control shRNA cassette in pGFP-V-RS Vector (Cat#TR30013) were purchased from Origene (Rockville, MD, USA). Lipofectamine™ stem transfection reagent (Cat#STEM00003) for cDNA cloning and shRNA, and lipofectamine RNAiMAX (Cat#13778030) for siRNA transfection were purchased from Life technologies (Carlsbad, CA, USA).

For SETD1A and WDR82 knock-down in hNSCs (StemPro®), sphere hNSCs were treated with WDR82 shRNA or its scrambled shRNA control (scrCtrl) using Lipofectamine™ stem transfection reagent or with 30nM hSETD1A and control siRNAs using lipofectamine RNAiMAX as per manufacturer’s instructions. For human SETD1A and WDR82 over-expression in normal HAs, HAs were plated in 6-well plates and transiently transfected with pET28-MHL or pET28-SETD1A-MHL, or with pcDNA3-WDR82-HA or pcDNA-HA using Lipofectamine™ stem transfection reagent as per manufacturer’s instructions. Transfected cells were subjected to sphere formation assays or lysed after 48h for quantitative real-time PCR, chromatin immunoprecipitation PCR and western blots.

### Neurosphere formation, cell proliferation and differentiation assays

Human StemPro® NSCs were harvested as single cells and plated on 12-well plates at 8.33 × 10^4^ cells/well and treated with 100ng/ml BMP4, or with buffer. Sphere formation was monitored daily and the number of spheres was counted 4 days after treatment. At 4 days the spheres were centrifuged and dissociated with StemPro® Accutase® into single cells. The cells were re-suspended in 1ml NSC SFM and counted with a hemocytometer. Sphere and cell numbers from triplicate wells were counted and analyzed. For sphere differentiation assays, human StemPro® NSCs were harvested as single cells and plated on 6-well plates at a density of 4.5 × 10^5^ cells/well. Sphere formation was monitored daily. On day 4 after plating, the spheres were treated with 100ng/ml BMP4, or buffer. Sphere differentiation was investigated daily for 7 days and cells were harvested for RNA extraction for PCR array assays.

For HA sphere formation assays, the cells were plated in 6-well plates and transfected with pET28-hSETD1A-MHL and pcDNA3-WDR82-HA using StemPro® NSC transfection reagent. After 48 hrs, the culture medium was replaced with Neurobasal medium supplement with N2, B27, EGF 20ng/ml and bFGF 20ng/ml. At day 4, sphere numbers from a minimum of ten fields in triplicate wells were counted and analyzed.

Sphere or cell numbers were averaged and graphed with Graphpad Prism 7.0 software (La Jolla, CA, USA). P values were calculated using 2-sided Student’s t test, with p<0.05 considered significant.

### Quantitative real-time PCR

Total RNA was isolated with the RNeasy Mini Kit (Qiagen, Valencia, CA, USA) from primary sphere cultures and spontaneously differentiated Human StemPro® NSCs, treated with BMP4 and from non-treated controls. cDNA was synthesized with qScript cDNA SuperMix (5 ×) (Quanta Biosciences, 95048-025) followed by real-time (RT) PCR with primers as indicated (Supplementary Table 1). To ensure accuracy, an internal reference reaction using GAPDH was performed on the same sample as used for the target gene. The results were standardized with the formula: ΔCT = CT_Ref_ − CT_Target_ and converted to fold change of target gene over reference gene (F = 2^-ΔCT^). Data from a minimum of 3 independent experiments were used to quantify gene expression. P values of less than 0.05 were considered statistically significant.

### Protein extraction and western blots

Total protein was extracted with RIPA buffer (Cell Signaling Technology, Cat#9806) with proteinase (Cell Signaling Technology, Beverly, MA, USA), phosphatase (Sigma) inhibitor cocktails and phenylmethylsulfonyl fluoride (PMSF, Roche). Total histone was extracted using a histone extraction kit (ab113476, Abcam) as per manufacturer’s instructions. Protein concentrations were quantified with the BCA Protein Assay Kit (Cat#23227, Thermo Fisher Scientific Inc.) with a Nanodrop ND-1000 (Thermo Fisher Scientific Inc.). Equal amounts of cell lysate or concentrated conditioned medium were resolved by sodium dodecyl sulfate– polyacrylamide gel electrophoresis and transferred to nitrocellulose membranes (Bio-Rad, Hercules, CA, USA). The membrane was blocked for 60min with 5% non-fat dry milk in Tris-buffered saline and Tween 20 (1:1000), followed by blotting with primary antibodies overnight at 4°C. Primary antibodies included: polyclonal anti-goat BMP4 (1:500), OCT3/4 (1:200), H3K4me3 (1:1000), H3 (1:2500), CYCLIN D1 (1:1000), β-ACTIN (1:3000), GAPDH (1:2000); and anti-human SETD1A (A300-289A, 1:1000) from Bethyl Laboratories. After washing with Tris-buffered saline and Tween 20, membranes were incubated for 1h at room temperature with horseradish peroxidase conjugated donkey anti-rabbit antibody (sc-2305, 1:5000), donkey anti-mouse antibody (sc-2306, 1:5000), or donkey anti-goat antibody (sc-2020, 1:5000) and signal was detected with enhanced chemiluminescence substrate (Bio-Rad).

### Chromatin immunoprecipitation (ChIP)

ChIP was performed according to the manufacturer’s instructions (Abcam). Briefly, 1 × 10^6^ cells were washed with ice cold PBS with proteinase inhibitor cocktail prior to fixation in 1% formaldehyde in PBS at room temperature for 10 min and quenched with 0.125 M glycine for 5 min. Cells were collected and re-suspended in 200μl FA-lysis buffer (50mM HEPES-KOH pH7.5, 140mM NaCl, 1mM EDTA pH8.0, 1% Triton X-100, 0.1% Sodium Deoxycholate, 0.1% SDS, 1mM PMSF, 1μg/mL leupeptin, 1μg/mL pepstatin) and disrupted using a 25 1/2 gauge syringe while on ice. The lysate was transferred into a 5ml conical tube and diluted with 1.3ml dilution buffer (0.01% SDS, 1.1% Triton X-100, 1.2mM EDTA pH 8.0, 16.7mM Tris-HCl pH 8.0, 167mM NaCl), sonicated and processed according to manufacturer’s protocol. Rabbit polyclonal H3K4me3 antibody (Cell Signaling Technology, Cat# 9727) and IgG from rabbit serum (Cell Signaling Technology, Cat# 2729) were used to collect immunoprecipitated DNA fragments. All DNA fragments were cleaned-up with GenElute™ PCR Clean-Up Kit (Sigma Aldrich Co., St. Louis, MO, USA) prior to PCR analysis. Human *OCT4* (Cell Signaling Technology, Cat#4641) and *CCND1* (Cell Signaling Technology, Cat#12531) and *NESTIN* (Cat# GPH1015041 (-) 01A, Qiagen, Germantown, MD, USA) promoter primers were used following real-time PCR analysis.

## RESULTS

### H3K4me3 is a stem marker for human NSCs

Slides from the region lining the lateral ventricle were prepared from two human fetal brain samples and immunostained for H3K4me3. H3K4me3 positive cells were enriched in the SVZ (Figure 1A), in agreement with previous findings from rodents [34] and baboons [13]. To determine if H3K4me3 and its methyltransferases hSETD1A and WDR82 were altered upon differentiation, RNA, total protein, and histones were extracted from spheres and from differentiated human NSCs (StemPro®). Real-time PCR and western blots both showed that hSETD1A and WDR82 expression was higher in sphere cultured cells compared to differentiated cells (Figure 1C and D). Other proteins included in the hSETD1A-COMPASS complex were not significantly changed (Supplementary Figure 1). H3K4me3 was readily detected in undifferentiated sphere-cultured hNSCs, but was undetectable in differentiated cells (Figure 1D).

**Figure 1.**
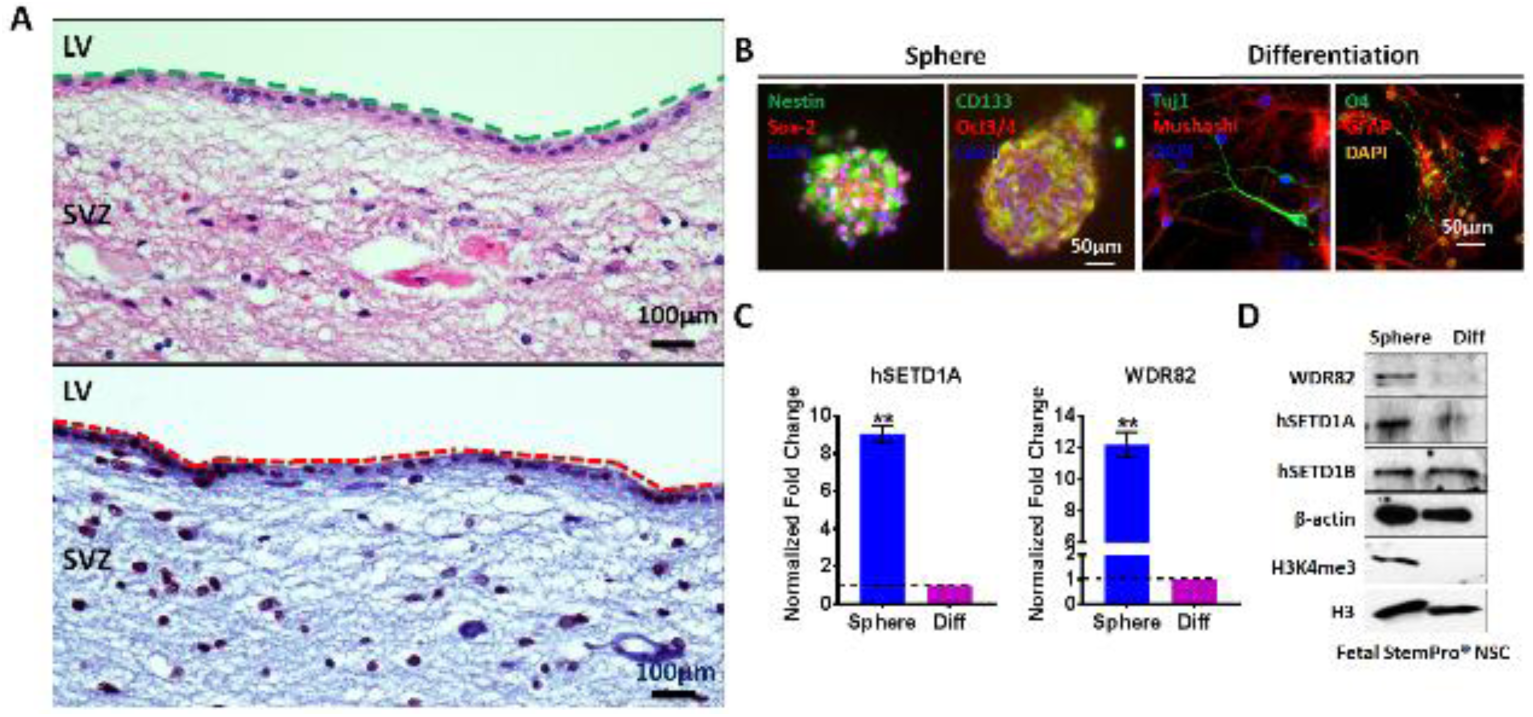
H3K4me3 is present in human NSCs undergoing self-renewal but not in differentiated cells. A. Gross anatomy of partial fetal SVZ architecture and localization of H3K4me3 positive cells. View of fetal brain in the area of the lateral ventricle (LV), showing the SVZ with H3K4me3 positive cells. Top panel: H&E staining (LV and SVZ are separated by a green dashed line). Bottom panel: immunostaining with H3K4me3 antibody (LV and SVZ are separated by a red dashed line). B. Immunofluorescence of human StemPro® NSC neurospheres positive for NESTIN, SOX2, CD133 and OCT4 and differentiated cells positive for MUSHASHI, GFAP, TUJ1 and O4. C. Real-time PCR showing hSETD1A and WDR82 expression in spheres and differentiated human StemPro® NSCs. D. Western blots using total protein extracts and histone from un- (UD) and -differentiated (Diff) cells. Error bars show the standard error of three independent experiments (** p<0.01).

These results indicate that H3K4me3 and the activity of its methyltransferases hSETD1A and WDR82 are associated with NSC self-renewal, but not with differentiated cells.

### H3K4me3 promoter occupancy determines hNSC fate

We examined the expression of OCT4 [35], a crucial stem cell self-renewal marker, and NESTIN, a neural stem/progenitor marker, for potential relationships with H3K4 methylation. Both were expressed in NSCs, with decreased expression in spontaneously differentiated cells, (Figure 2 A-E). Using H3K4me3 ChIP coupled with real-time PCR (Figure 2F) we determined that H3K4me3 is enriched at OCT4 and NESTIN promoters in undifferentiated hNSCs relative to their differentiated derivatives (Figure 2F).

**Figure 2.**
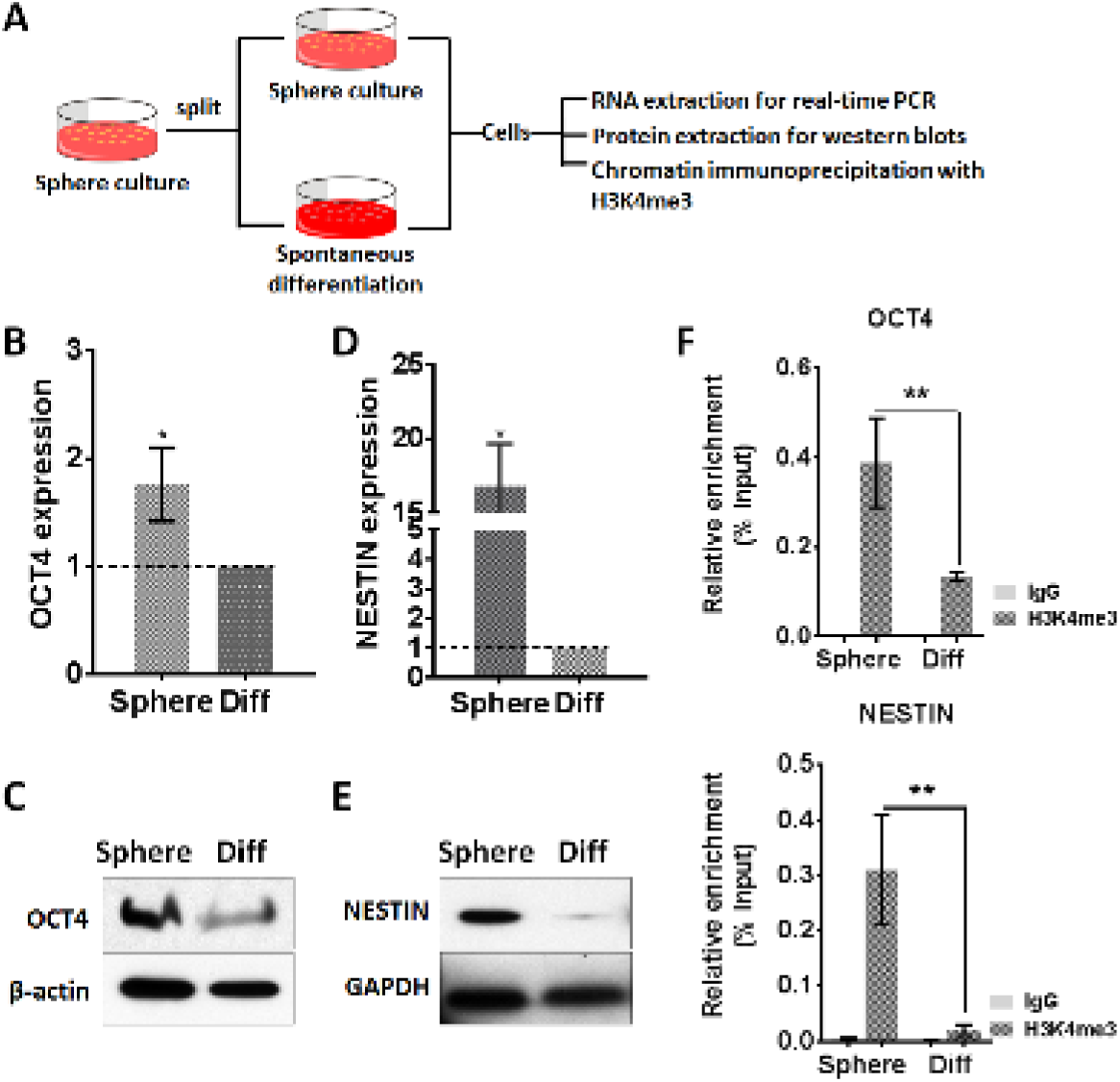
H3K4me3 is present at promoters of genes associated with NSC self-renewal. A. Experimental design for sample collection. B-E Real-time PCR (B and D) and western blots (C and E) showing OCT4 and NESTIN mRNA and protein in undifferentiated spheres (Sphere) and spontaneously differentiated (Diff) human StemPro® NSCs. F. Chromatin-immunoprecipitation (ChIP) with rabbit IgG and H3K4me3 and detected with real-time PCR using promoter primers. Error bars show the standard error of three independent experiments (* p<0.05, ** p<0.01).

### BMP4 decreases H3K4me3 at CCND1 promoter in differentiated hNSCs

To determine whether BMP4 inhibits human StemPro® NSCs proliferation through reduction of H3K4me3 at the CYCLIN D1 gene, hNSC neurospheres were cultured in 100ng/ml BMP4 (Figure 3A), which reduced both the number of neurospheres (diameter >250μm) (Figure 3B and C) and the overall number of cells (Figure 3D). BMP4 reduced CYCLIN D1 expression (Figure 3F and G) and CYCLIN D1 promoter H3K4me3 occupancy (Figure 3H).

**Figure 3.**
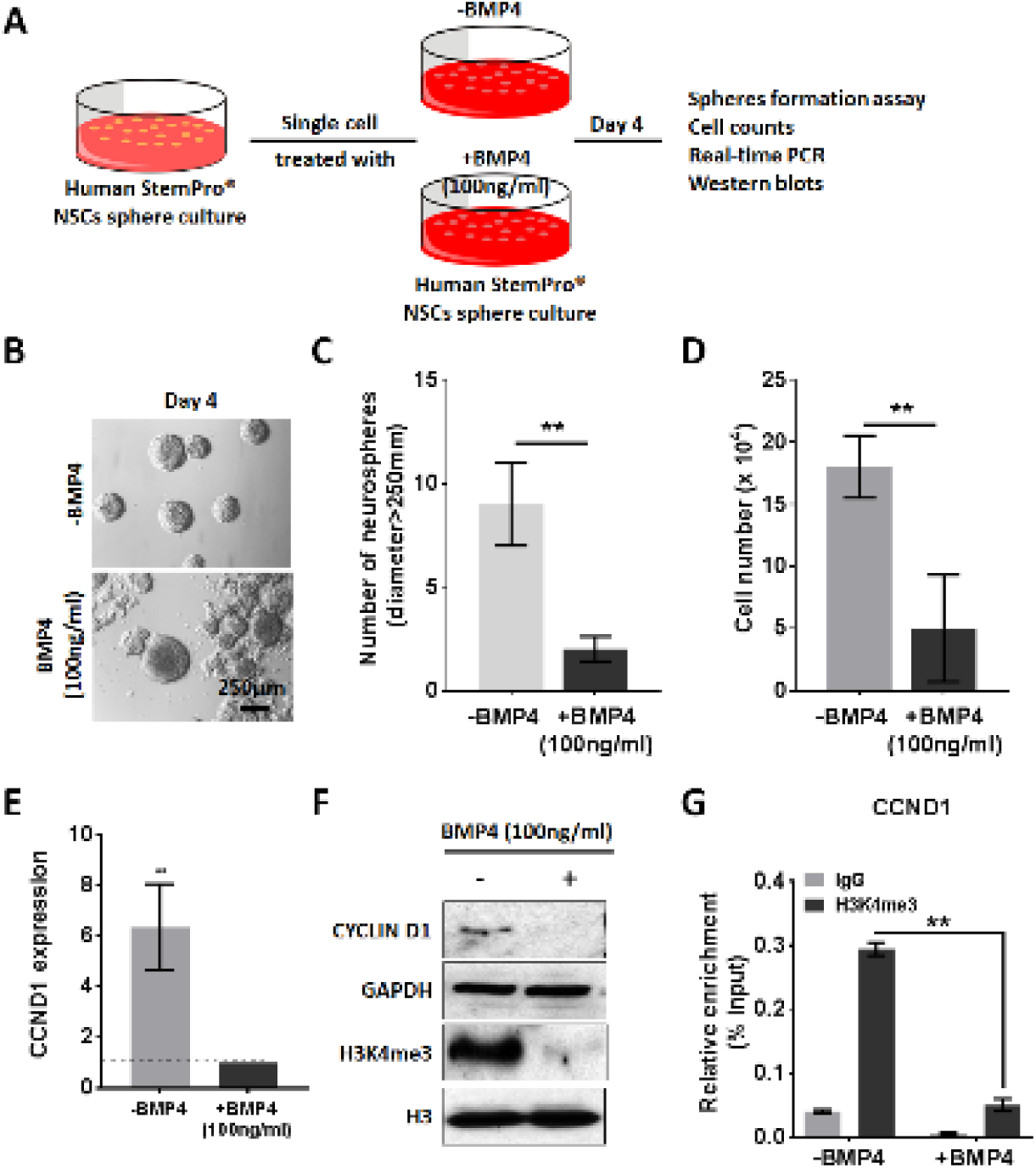
BMP4 inhibits human neurosphere formation and decreases H3K4me3 at the CYCLIN D1 promoter. A. Experimental design. B, C and D. Representative images showing sphere morphology (B), number of neurospheres (diameter > 250μm) (C) and cell numbers (D) at day 4. E. Western blots showing H3K4me3 levels. E and F. Real-time PCR (E) and western blot (F) for CYCLIN D1. G. Chromatin-immunoprecipitation (ChIP) with rabbit IgG and H3K4me3 combined with real-time PCR to detect CYCLIN D1 promoter. Error bars show the standard error of three independent experiments (** p<0.01).

### BMP4 reduces H3K4me3 at the NESTIN promoter and promotes hNSC differentiation

We monitored differentiation of neurospheres from hNSCs following treatment with 100ng/ml BMP4 vs untreated controls (Figure 4A). By the fourth day nearly all neurospheres treated with BMP4 had attached and migrated (Figure 4B). Additionally, H3K4me3 markedly decreased in the treatment group compared to the controls (Figure 4C). NESTIN expression, examined by real-time PCR (Figure 4D) and western blot (Figure 4E), showed significant decreases with BMP4 treatment, as did levels of H3K4me3 at the NESTIN promoter (Figure 4F).

**Figure 4.**
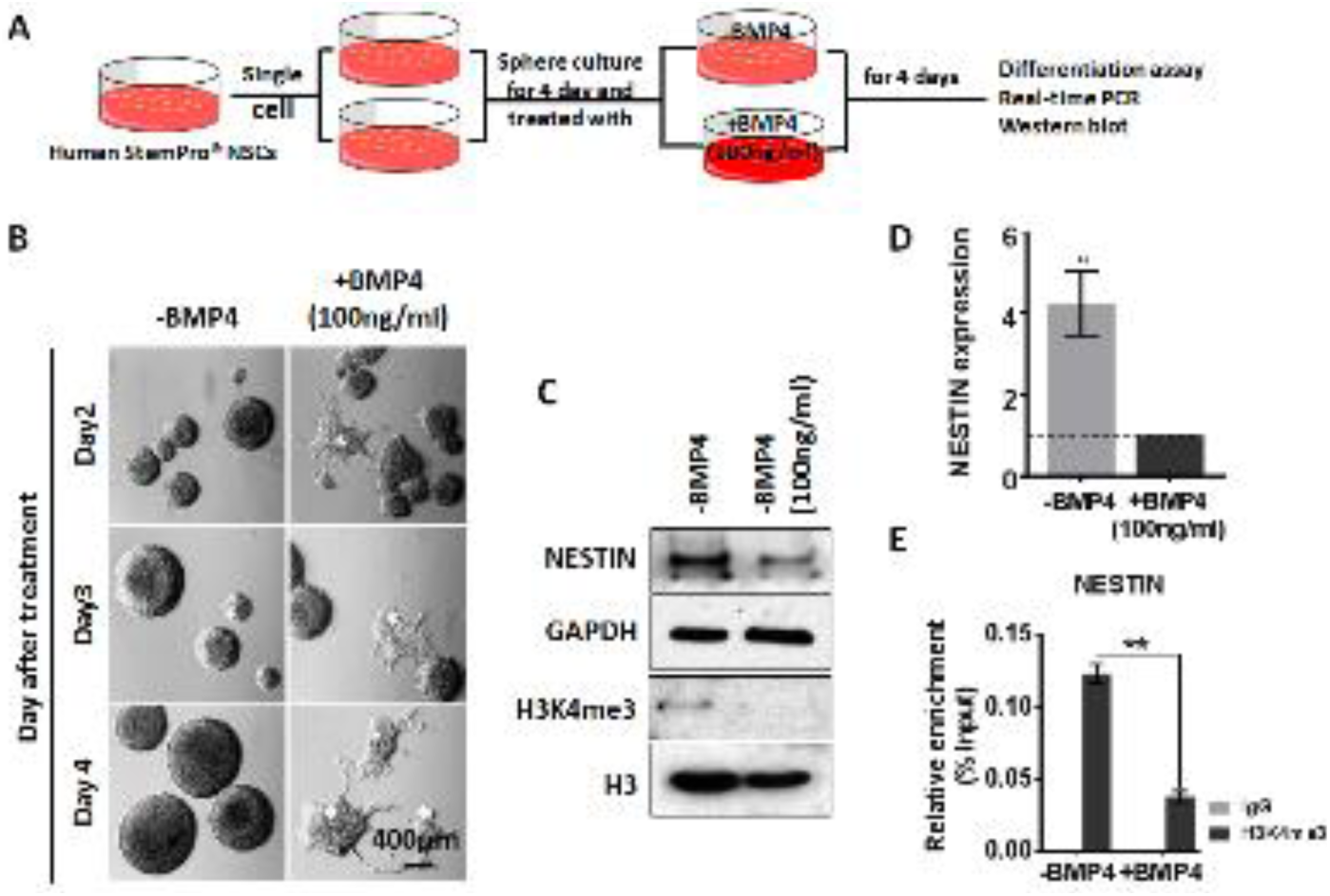
BMP4 drives human neurosphere differentiation with reduction of NESTIN promoter H3K4me3 occupancy. A. Experimental design. B. Representative images showing sphere morphology +BMP4 100ng/ml, in comparison to control (-BMP4). C. Western blots using total and histone extracts to detect NESTIN and H3K4me3, GAPDH and H3 as controls. D. Real-time PCR shows expression of NESTIN. E. Chromatin-immunoprecipitation (ChIP) with rabbit IgG and H3K4me3 combined with real-time PCR using NESTIN promoter primers. Error bars show the standard error of three independent experiments. (** p<0.01)

### BMP4 reduces SETD1A and WDR82 expression, with a corresponding reduction in cellular H3K4me3

BMP4 reduced levels of WDR82 and human SETD1A levels (Figure 5A). To determine the biologic effects in NSCs of reduced SETD1A and WDR82, hNSCs were incubated with SETD1A siRNA or transduced with lentivirus expressing WDR82 shRNA. Both treatments decreased sphere numbers in comparison to controls (Figure 5B). Target transcripts as well as transcripts associated with stemness (OCT4 and NESTIN) and proliferation (CCND1) also decreased (Figure 5C and D). H3K4me3 chromatin immunoprecipitation coupled with real-time PCR revealed that these transcript reductions occurred in conjunction with decreased H3K4me3 promoter occupancy (Figure 5E).

**Figure 5.**
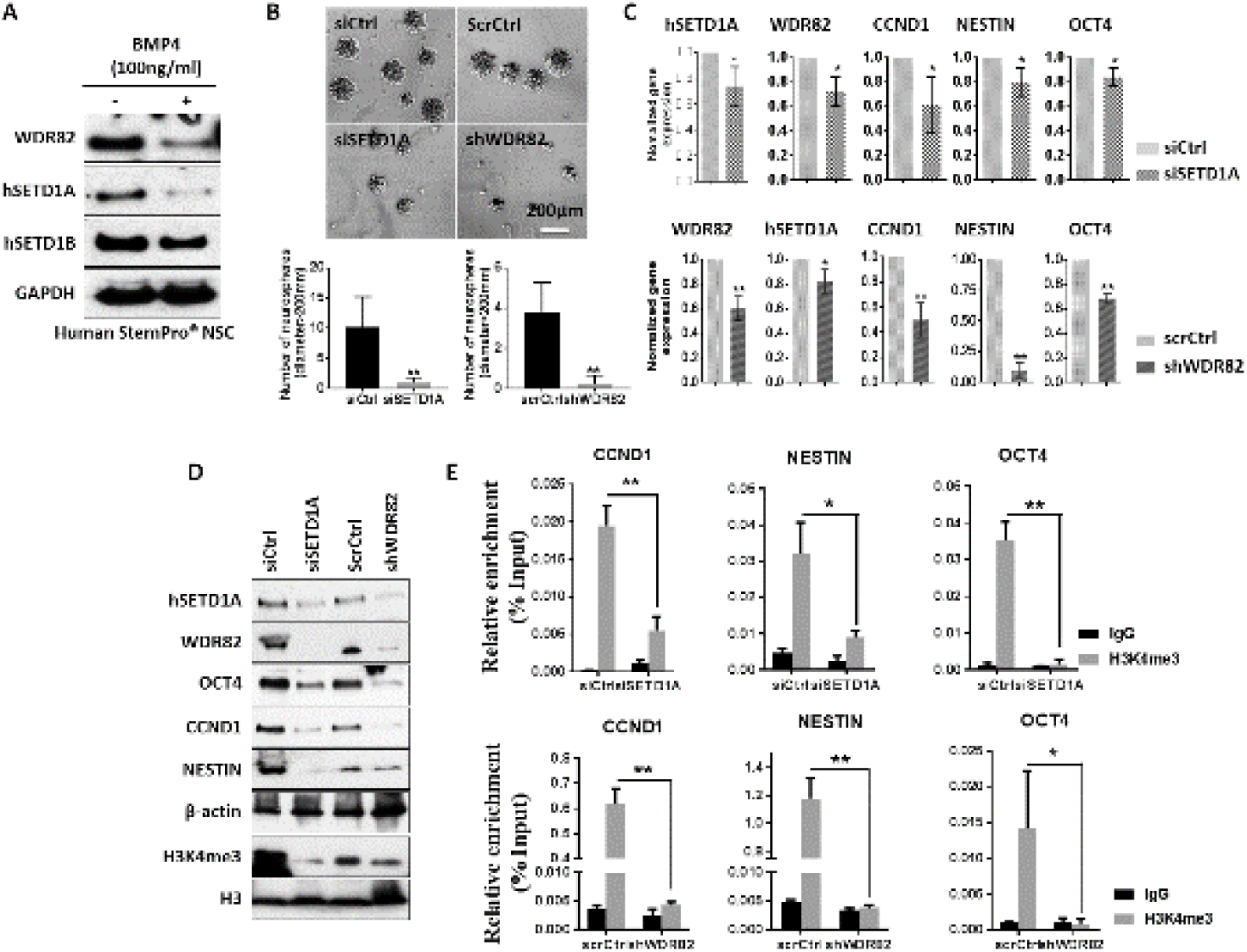
BMP4 reduces global H3K4me3 promoter occupancy through reduction of its methyltransferases WDR82 and hSETD1A. A. Western blots shows WDR82 and human SETD1A/B expression following 100ng/ml BMP4 treatment. B. Representative images and quantitative graphs showing sphere formation in human StemPro® NSCs following treatment with siRNA for hSETD1A (siSETD1A) or shRNA for WDR82 (shWDR82) vectors versus controls (control siRNA, siCtrl and scrambled shRNA, scrCtrl). C and D. Real-time PCR (C) and western blots (D) showing expression of hSETD1A, WDR82, OCT4, CCND1 and NESTIN, following treatments as in B. E. Real-time PCR using DNA from ChIP with rabbit IgG and H3K4me3 and detected with promoter primers for OCT4, CCND1 and NESTIN following treatment with siSETD1A, shWDR82 or siCtrl, scrCtrl, in StemPro® NSCs. Error bars show the standard error of three independent experiments. (* p<0.05, ** p<0.01)

### Ectopic expression of SETD1A and WDR82 increases expression of stemness and proliferation genes in normal human astrocytes

We investigated whether increasing expression of WDR82 or hSETD1A in human astrocytes (HAs) promotes cellular dedifferentiation. When astrocytes were transfected with WDR82 or hSETD1A cDNA, and cultured in hNSC medium for 4 days, sphere numbers increased (Figure 6A), as did expression of OCT4, CCND1 and NESTIN, indicated by real-time PCR and western blot analysis. (Figure 6 B and C). H3K4me3 promoter occupancy at these genes also increased (Figure 6D).

**Figure 6.**
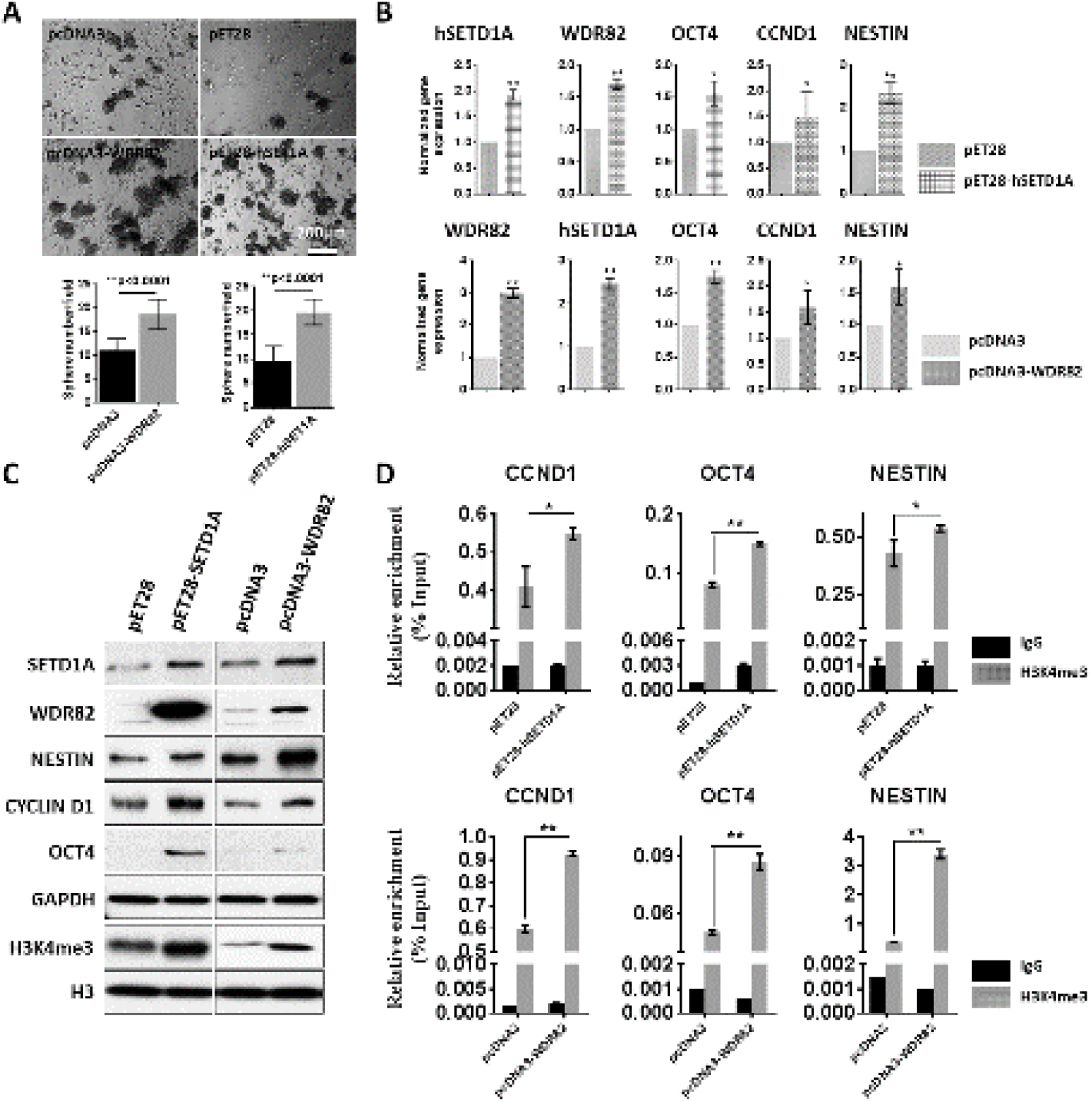
Ectopic expression of human SETD1A and WDR82 increases global H3K4me3 promoter occupancy in normal human astrocytes (HAs). A. Representative images and quantitative graphs show sphere formation from HAs following transfection with WDR82 (pcDNA3-WDR82) or human SETD1A (pET28-hSETD1A-MHL) expression plasmids, in comparison to controls (pcDNA3 and pET28- MHL). B and C. Real-time PCR (B) and western blots (C) show expression of hSETD1A, WDR82, OCT4, CCND1 and NESTIN. D. ChIP with rabbit IgG and H3K4me3 combined with real-time PCR using promoter primers for OCT4, CCND1 and NESTIN following HA transfection. Error bars show the standard deviation of three independent experiments. (* p<0.05, ** p<0.01)

## DISSCUSSION

BMP4 is critical for nervous system development due to its influence over several biological processes, ranging from dorsoventral patterning of the embryonic neural tube to proliferation, apoptosis, neurogenesis, and gliogenesis at different stages and locations [36-38]. BMP4 also plays important roles in the regulation of neural progenitor cell fate and promotes neuronal precursor differentiation in the spinal cord [39] and cortex [40]. Here we found that cellular H3K4me3 and its methyltransferases hSETD1A and WDR82 decrease upon BMP4-induced hNSC differentiation. This differentiation is accompanied by decreased expression of stem cell signature proteins OCT4 and NESTIN, in conjunction with decreased H3K4me3 at their promoters. We also showed that BMP4 reduces proliferation of human NSCs and inhibits neurosphere formation, this may be through decreased CYCLIN D1 promoter H3K4me3 occupancy, and thus CYCLIN D1 expression. Conversely, increased expression of WDR82 and hSETD1A in normal HAs increased global H3K4me3 levels and promoter occupancy at genes associated with stemness, proliferation and differentiation of hNSCs. Collectively these findings clearly demonstrate that hSETD1A and WDR82 are critical for maintaining neural stem cell populations, and decreasing in their expression promotes cellular differentiation.

Recent studies have shown histone PTMs are essential in hNSC self-renewal and differentiation. For example, H3K4me3 is present in undifferentiated progenitor cells within the SVZ and it is associated with distinct neurogenesis networks and pathways [13]. H3K4me3 is also required to ensure transcriptional precision at key genes during neural progenitor cell differentiation [20]. In the present study, we show that BMP4 changes H3K4me3 levels at specific loci including those involved with stemness (OCT4 and NESTIN) and proliferation CCND1 (Figures 2-4).

Mammalian SETD1A/B-COMPASS are responsible for H3K4me3 [41], with SETD1A most relevant in the development and maintenance of hNSCs [42, 43]. In silico analysis results from several published databases (GSE55379, GSE53404, and GSE43382) show high levels of SETD1A in human NSCs, with decreased levels in differentiated cells (Supplementary Figure 2A). In this study we show that SETD1A is high in human NSCs and decreases in differentiated cells (Figure 1C and D), in agreement with previous findings.

Our results show that expression of WDR82, a subunit of SETD1A-COMPASS, is also high in human NSCs, and that it decreases in differentiated cells (Figure 1C and D). WDR82 mediates alterations in H3K4me3 levels and in so doing influences hNSC proliferation and apoptosis during normal embryonic growth and development [44]. Analysis of results from databases GSE15209 and GSE43382 show that WDR82 is high in human NSCs, but decreases in differentiated brain cortex (GSE15209) and differentiated human NSCs (GSE43382) (Supplementary Figure 2B). This is consistent with our results and highlights WDR82 involvement in regulating hNSC stemness and differentiation.

SETD1A acts through several mechanisms to bring about intracellular changes. Studies have shown that it modulates cell cycle progression through induction of a miRNA network to suppress P53 in human cancers [45, 46], independent of H3K4me3. A non-catalytic domain, “FLOS” (functional location on SETD1A), interacts with CYCLIN K and regulates DNA damage response gene expression [47]. In the present study, we found reduction of WDR82 decreased hSETD1A expression in hNSCs (Figure 5C and D), while ectopic expression of WDR82 markedly increased SETD1A (Figure 6B and C) in normal HAs, consistent with findings from other groups [48, 49]. Conversely, reduction of hSETD1A decreased hNSC WDR82 expression (Figure 5C and D), whereas induction of SETD1A markedly increased WDR82 in normal HAs (Figure 6B and C). WDR82 interacts with an RNA recognition motif (RRM) [48, 49], consisting of 95 highly conserved amino acids, at the N-terminal of hSETD1A protein (Supplementary Figure 3). Our findings indicate that WDR82 and hSETD1A control cellular H3K4me3 in hNSCs and suggest that BMP4 promotes hNSC differentiation through an epigenetic mechanism.

## CONCLUSION

In this work, we have shown that H3K4me3 promoter occupancy mediated by hSETD1A and WDR82 is critical for hNSC fate. BMP4 decreases hSETD1A and WDR82, resulting in the reduction of H3K4me3 at key genes such as OCT4, NESTIN and CCND1 to inhibit proliferation and promote differentiation of hNSCs (Figure 7). These results extend our understanding of the role of histone PTMs in normal CNS development.

**Figure 7.**
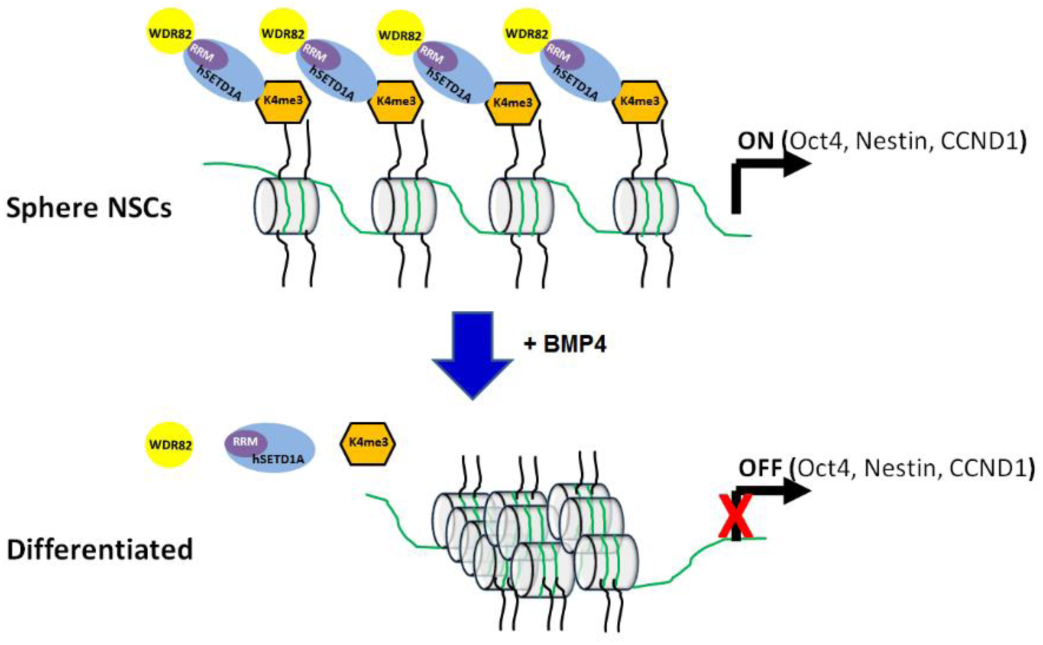
Schematic diagram showing that BMP4 decreases levels of H3K4me3 mediated by hSETD1A-WDR82 at promoters of key regulators to promote differentiation and inhibit proliferation of human neural stem cells (NSCs).

## ACKNOELEDGEMENTS

This project was partially supported by National Cancer Institute SPORE grant (P50CA221747-01A1) Career Enhancement Award and by the Rory David Deutsch Foundation, the Surgical Neuro-Oncology Research Fund of Ann & Robert H. Lurie Children’s Hospital (A&RLCH) of Chicago, and the Dr. Ralph and Marian C. Falk Medical Research Trust.

We appreciate generosity of Dr. David G. Skalnik at Indiana University School of Medicine to provide rabbit polyclonal antibody against WDR82 and pcDNA3-WDR82-HA plasmid. We also thank Drs. Kunliang Guan and Cheryl Arrowsmith who created pcDNA3-HA and pET28-SETD1A-MHL and its control pET28-MHL plasmids, respectively and donated to Addgene.

