## Supplementary file for "Bone morphogenetic protein 4 reduces global H3K4me3 to inhibit proliferation and promote differentiation of human neural stem cells"

**RUNNING TITLE: BMP4 reduces H3K4me3 in human NSCs**

Sonali Nayak,^1^ Benjamin Best,^1^ Emily Hayes,^1^ Yuping D Li,^1^ Gordan Grahovac,^1^ Nitin R Wadhwani,^2^ Barbara Mania-Farnell,^3^ Rintaro Hashizume,^4^ Chandra S Mayanil,^1, 4, 5^ John A Kessler,^6^ Charles David James,^4^ Tadanori Tomita,^1, 4^ Guifa Xi,^1, 4, 5^

^1^Falk Brain Tumor Center, Division of Pediatric Neurosurgery, ^2^Department of Pathology, Ann & Robert H. Lurie Children’s Hospital of Chicago, Northwestern University Feinberg School of Medicine, Chicago, IL 60611

^3^Department of Biological Sciences, Purdue University Northwest, Hammond, IN, 46323

^4^ Department of Neurological Surgery, Northwestern University Feinberg School of Medicine, Chicago, IL 60611

^5^ Developmental Biology Program, Stanley Manne Research Institute, Ann & Robert H. Lurie Children’s Hospital of Chicago, Northwestern University Feinberg School of Medicine, Chicago, IL 60611

^6^Department of Neurology, Northwestern Memorial Hospital, Northwestern University Feinberg School of Medicine, Chicago, IL 60611

**Corresponding author:**

Guifa Xi, MD, PhD

Affiliation: Division of Pediatric Neurosurgery, Ann & Robert H. Lurie Children’s Hospital of Chicago, Northwestern University Feinberg School of Medicine

Address: 225 E Chicago Ave, PO Box #28, Chicago, IL 60611

**Supplementary Table 1** List of Real-time PCR primers used in this study

| **Name** | **Forward Primer (5’-3’)** | **Reverse Primer (5’-3’)** |
| --- | --- | --- |
| OCT4  NESTIN  CCND1  SETD1A | CAGGAGATATGCAAAGCAGAAAC  GGCAGCG TTGGAACAGAGGT  TTCGGGATGATTGGAATAGC  CGGAAGAAGAAGCTCCGATTT | GGCACTGCAGGAACAAATTC  CATCTTGAGGTGCGCCAGCT  TGTGAGCTGGCTTCATTGAG  ACCCTATTGTGCCTCAGTTTC |
| WDR82 | AGAGCAGAGAGAGGAGTATGTC | ACCCTATTGTGCCTCAGTTTC |
| GAPDH | TGACATCAAGAAGGTGA | TCCACCACCCTGTTGCTGTA |

**Supplementary Figures**

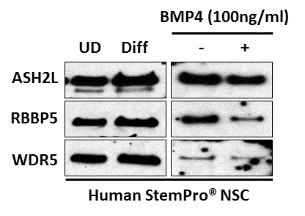

**Supplementary Figure 1**. Immunoblots showing expression of human SETD1A subunits ASH2L, RBBP5, and WDR5 in protein extracted from un- (UD) and differentiated (Diff), non- (-) and BMP4 (100ng/ml) (+) treated human StemPro® NSCs.

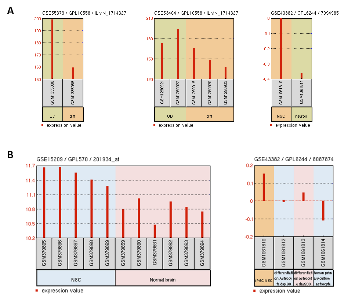

**Supplementary Figure 2.** Expression patterns of SETD1A (A) and WDR82 (B) in human neural stem cells (NSCs) and differentiated cells, neurons or astrocytes, derived from hNSCs through in silico analysis of indicated datasets. UD: undifferentiated; Diff: differentiated; iPSC: induced pluripotency stem cell.

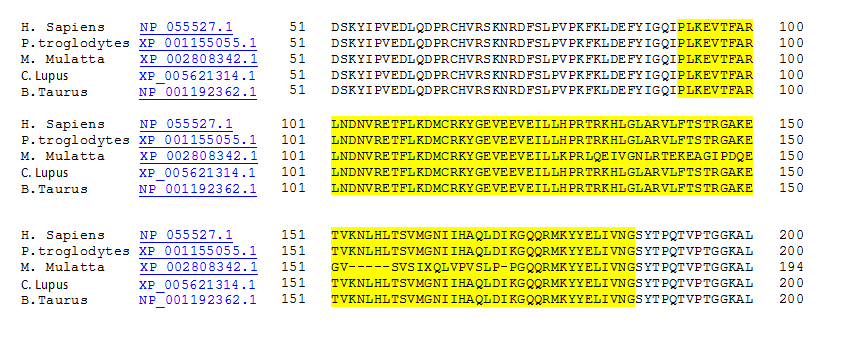

**Supplementary Figure 3.** The RNA recognition motif (RRM) (yellow highlighted) at the N-terminal of SETD1A, is a highly conserved site for WDR82 interaction.
